# Sequestration of clock proteins into repressive nuclear condensates orchestrates circadian gene repression

**DOI:** 10.1101/2025.11.03.686412

**Authors:** Qianqian Chen, Ye Yuan, Dunham Clark, Swathi Yadlapalli

**Author notes:** Corresponding author: Swathi Yadlapalli.

## Abstract

Circadian clocks orchestrate ∼24-hour cycles in gene expression, behavior, and physiology across most organisms^1^. Our recent study has revealed a striking spatial organization in the nucleus during the repression phase: core clock proteins such as *Drosophila* PERIOD (PER) are organized into distinct nuclear foci close to the inner nuclear envelope of clock neurons, and clock-regulated genes are similarly positioned in the nucleus^2^. However, the functional relevance of this subnuclear organization is unknown. Here, we investigate how the spatial organization of clock proteins and chromatin regulates circadian gene repression. Given that PER partners with TIMELESS (TIM) to enact transcriptional repression, we first investigated whether TIM is also a component of these nuclear foci. Using CRISPR-Cas9 to endogenously tag TIM with mNeonGreen, we performed high-resolution live imaging in *Drosophila* clock neurons. We found that TIM forms nuclear foci during the repression phase that co-localize with PER condensates, whereas TIM remains diffuse in the cytoplasm of *per^01^* null mutants. To probe the spatial relationship between these condensates and clock-regulated genes, we combined protein imaging with fluorescence in situ hybridization (FISH) which revealed that PER/TIM condensates were positioned adjacent to, but did not overlap with, clock gene loci during the repression phase. These results suggest that core clock proteins are spatially sequestered into repressive condensates away from chromatin, providing a new framework for understanding how nuclear architecture and phase separation together orchestrate rhythmic gene expression.

## INTRODUCTION

Circadian clocks are cell-autonomous timekeepers regulating ∼24-hour rhythms in gene expression, behavior, and physiology across eukaryotes^1^. They function through a highly conserved transcriptional-translational feedback loop. In *Drosophila*, transcriptional activators CLOCK (CLK) and CYCLE (CYC) induce clock genes, including those encoding repressors such as PERIOD (PER) and TIMELESS (TIM). These repressors accumulate, translocate into the nucleus after a delay, and inhibit the activators, suppressing their own transcription and that of other clock-regulated genes (Figure 1A). While past studies employing genetic screens and biochemical assays have identified key clock components and genetic circuits underlying circadian regulation^1, 3-5^, how these are spatially organized and dynamically regulated at the subcellular level remains poorly understood. To address this, we use *Drosophila melanogaster*, whose molecular clock mirrors humans’^1^, leveraging its well-characterized network of ∼150 clock neurons (Figure 1B).

**Figure 1.**
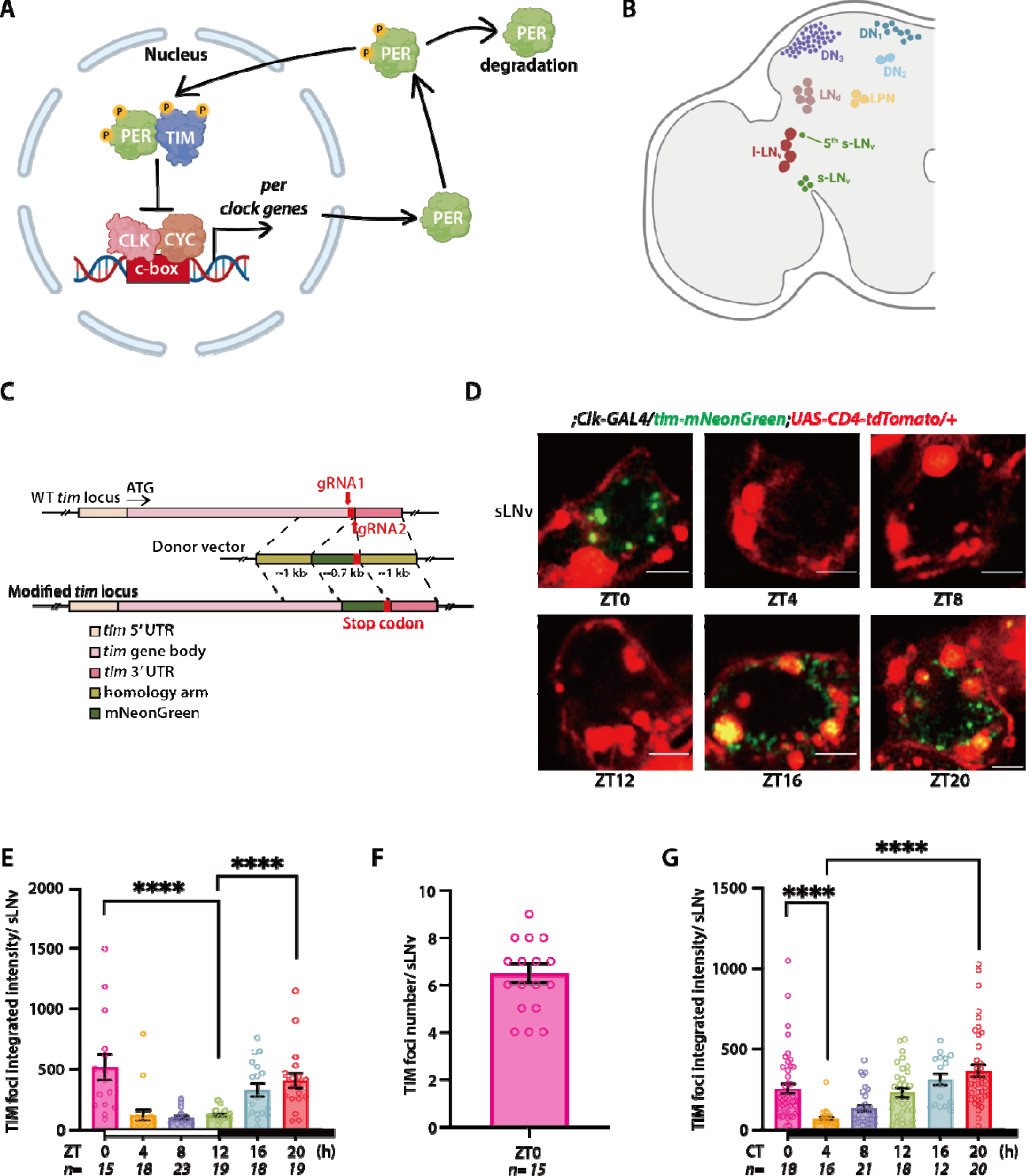
TIM protein is organized into a few discrete foci during the circadian repression phase. (A) Schematic of the core molecular clock in *Drosophila* clock neurons. During the activation phase, the CLK protein complex binds to the E-box sequence of the hundreds of clock-regulated genes and drives their transcription, including *per* and *tim*. During the repression phase, the PER protein enters the nucleus with TIM and inhibits CLK transcription factor activity, silencing its own expression. (B) Schematic of the *Drosophila* circadian clock network, with the major classes of clock neurons labeled. (C) Schematic of CRISPR/Cas9 genome editing to generate *Tim-mNeonGreen* flies. (D) Representative images of TIM-mNeonGreen (green) in s-LNv neurons at different zeitgeber times (ZT0, ZT4, ZT8, ZT16, ZT20). Scale bars-2µm. CD4tdTom (red) is used to label the membranes in the cytoplasm of clock neurons. (E) Quantification of TIM protein fluorescence intensity per s-LNv over the light-dark cycle. (F) Quantification of number of TIM foci per s-LNv at ZT0. (G) Quantification of TIM protein fluorescence intensity per s-LNv in the third day of constant darkness. Flies are entrained to LD cycles for 5 days and released into DD. CT refers to circadian time. The statistical test used was a Kruskal-Wallis test (E, F, and G). *P < 0.05, ****P < 0.0001. Individual data points, mean, and SEM are shown. ‘n’ refers to the number of neurons. Scale bars-2µm.

Traditionally, clock proteins, genes, and mRNAs were thought to diffuse freely, encountering each other by random collisions^1^. However, emerging evidence challenges this view and suggests that spatiotemporal organization of clock proteins and genes is critical for circadian regulation^2, 6-11^. We found that Drosophila clock proteins are organized into dynamic, membrane-less condensates near the inner nuclear envelope specifically during the repression phase^2^. Additionally, DNA-FISH revealed that core clock gene loci, such as *per* and *tim*, are also positioned in close proximity to the nuclear envelope^2^. Similar patterns have begun to emerge across species: clock protein nuclear bodies have been observed in mammalian cells^8^ and the fungus Neurospora^9^, and clock-regulated genes are positioned close to the nuclear periphery in human U2OS cells^12^. Because many clock proteins contain large intrinsically disordered regions (IDRs)^13, 14^, they may undergo phase transitions that organize circadian regulation through condensate dynamics^15^. How broadly conserved this phenomenon is, and whether other core clock proteins (e.g., TIM, PER’s canonical partner) also form such condensates, remain open questions. Furthermore, the precise relationship between these clock protein condensates and chromatin at the nuclear envelope, particularly how their spatial positioning influences transcriptional repression, has not yet been explored in any model system.

To directly address these questions, we first asked whether other core clock proteins, e.g., TIM, co-localizes with PER nuclear condensates during the repressor phase. We then combined protein imaging with fluorescence in situ hybridization (FISH) to simultaneously visualize clock protein condensates and gene loci. Our findings demonstrate that the PER/TIM repressor complex is organized into distinct nuclear condensates, which are spatially segregated from their target genes. This discovery highlights the critical role of biomolecular condensates in circadian regulation: by precisely controlling the location and timing of clock protein interactions with target genes, these membrane-less condensates ensure robust and temporally accurate gene expression. This insight reveals a novel mechanism through which spatial organization governs circadian rhythms, offering broader perspectives on how nuclear architecture fine-tunes gene expression programs.

## RESULTS AND DISCUSSION

### Timeless protein is organized into discrete nuclear foci during the repression phase

In our previous work, we demonstrated that the endogenous fluorescently tagged PERIOD (PER) protein forms discrete nuclear foci during the circadian repression phase^2^. Because PER interacts with TIMELESS (TIM) to regulate transcription through the core negative feedback loop (Figure 1A), we investigated whether TIM exhibits a similar subcellular organization. Towards this goal, we used CRISPR/Cas9 to fuse mNeonGreen to the terminal exon of the endogenous *tim* gene (Figure 1C). mNeonGreen is a bright, monomeric fluorescent protein (∼3-5× brighter than EGFP) with a ∼3-fold faster maturation rate, making it well suited for live imaging^16^. We confirmed that the TIM-mNeonGreen (TIM-mNG) fusion protein is fully functional *in vivo*: flies carrying the *tim-mNeonGreen* allele displayed wild-type locomotor activity rhythms under both 12 h light-dark (LD12:12) cycles and constant darkness (DD), with a ∼24 hour free-running period (Figure S1A, B, D). In these flies, *per* mRNA levels also oscillated with normal amplitude and phase (Figure S1E). Moreover, a single copy of the *tim-mNeonGreen* allele rescued rhythmicity in *tim^01^* null mutants (Figure S1C, D), demonstrating that the fusion protein supports a fully functional molecular clock.

To track TIM localization and dynamics, we entrained TIM-mNG flies to LD12:12 cycles (ZT0-lights on, ZT12-lights off) and imaged clock neurons from live brains every 4 hours throughout the circadian cycle using Airyscan high-resolution confocal microscopy. We first focused on the small ventrolateral neurons (s-LNvs), the master pacemaker neurons that express pigment-dispersing factor (PDF). TIM levels exhibited robust oscillations over the circadian cycle: TIM was absent during the day, diffusely cytoplasmic in the early night, became nuclear by ZT20, and formed discrete, dynamic nuclear foci at ZT0 (Figure 1D, E). The presence of discrete nuclear foci at ZT0 was confirmed using an NLS-BFP marker to label the nuclei of clock neurons (Figure 2A). TIM foci were enriched near the nuclear periphery, with an average distance from the nuclear envelope of <0.5 μm (Figure S2D). In contrast to PER, whose nuclear foci persist until ZT6^2^, TIM foci rapidly degraded upon light exposure (Figure 1D). This light-dependent degradation is mediated by the photoreceptor Cryptochrome (Cry), which drives clock resetting through TIM degradation^17^. Similar temporal and spatial patterns were observed in the large ventrolateral neurons (l-LNvs) (Figure S2). Under constant darkness, TIM fluorescence intensity continued to oscillate with a ∼24-hour periodicity, although degradation occurred more slowly than under LD conditions (Figure 1G). The formation of discrete TIM nuclear foci contrasts with earlier immunofluorescence studies of paraformaldehyde-fixed tissue, which reported a diffuse nuclear TIM distribution. This discrepancy likely arises because condensates disassemble upon fixation^15^, a phenomenon we previously observed for PER condensates^2^, explaining why these structures were not detected in earlier work. Together, these findings reveal that TIM, like PER, is organized into discrete nuclear foci during the circadian repression phase.

**Figure 2.**
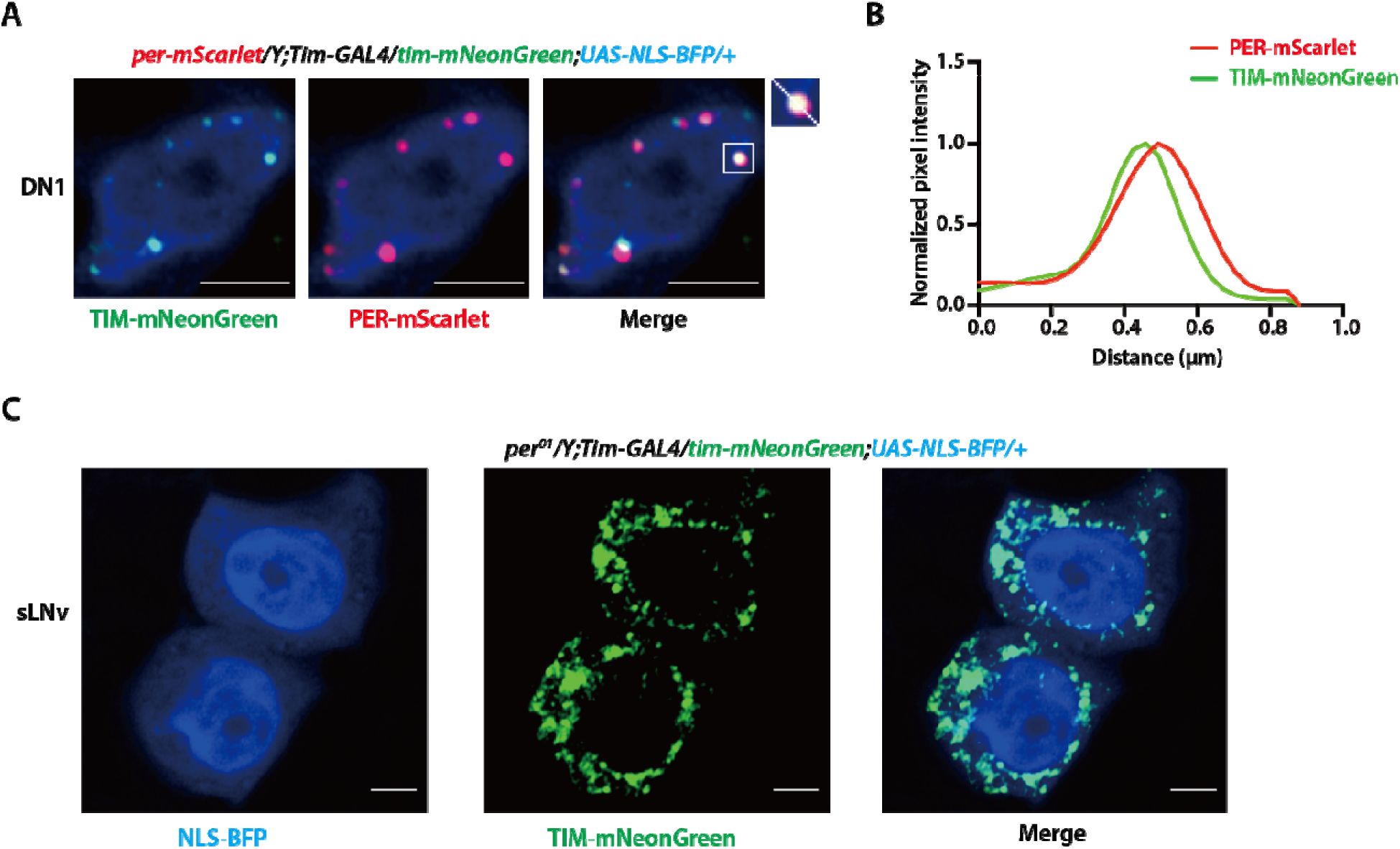
TIM and PER nuclear foci are colocalized during the repression phase. (A) Representative images of TIM-mNeonGreen protein and PER-mScarlet protein in DN1 neuron at ZT0. Nuclei were labeled with NLS-BFP (blue). Inset shows magnified view of a representative colocalization event. Scale bars-2µm. (B) Line profile analysis of the boxed region in (A), generated with ImageJ. Normalized pixel intensity of TIM-mNeonGreen (green) and PER-mScarlet (red) along the indicated line demonstrates overlapping peaks, indicating colocalization. (C) Representative images of TIM-mNeonGreen in sLNv neurons from *per^01^* null mutant flies. Nuclei were labeled with NLS-BFP (blue), and TIM was visualized as TIM-mNeonGreen (green). In the absence of PER, TIM failed to accumulate in the nucleus and remained cytoplasmic. Scale bars-2µm.

### PER and TIM foci co-localize at the nuclear periphery during the repression phase

Previous biochemical studies have established that the core clock proteins PERIOD (PER) and TIMELESS (TIM) form a PER/TIM complex during the repression phase to inhibit transcriptional activation by CLOCK/CYCLE (CLK/CYC). To determine whether PER and TIM foci co-localize in vivo, we used CRISPR-Cas9 genome editing to endogenously tag the per gene at its C-terminus with mScarlet-I, a bright, fast-maturing monomeric red fluorescent protein^18^. Because we had previously shown that *per-mNeonGreen* is fully functional^2^, we used the same guide RNAs to generate the *per-mScarlet-I* transgenic fly line and confirmed its full functionality in vivo through behavioral assays.

For live imaging, we generated flies with the genotype *per-mScarlet-I; tim-mNeonGreen; Tim>NLS-TagBFP*, which simultaneously express PER-mScarlet and TIM-mNeonGreen, with clock neuron nuclei labeled by nuclear-localized TagBFP. (NLS is a short peptide sequence that directs proteins into the nucleus.) We focused on the dorsal neuron 1 (DN1) subset, as these clock neurons are located near the brain surface, facilitating visualization of mScarlet fluorescence in live preparations. We observed strong co-localization of PER-mScarlet and TIM-mNeonGreen foci at the nuclear periphery during the repression phase (Figure 2A, B). This provides direct evidence that the PER/TIM repressive complex is organized into discrete nuclear foci. Importantly, we previously demonstrated that PER also sequesters the activator CLK into these same foci. Taken together, these results provide a direct visualization of the core repression mechanism, showing that the key repressors (PER and TIM) and their target activator (CLK) are compartmentalized within the same dynamic nuclear foci.

Because PER/TIM heterodimerization is required for their nuclear translocation^19^, we next examined TIM localization in *per*_*¹* null mutants at ZT0, a time when TIM forms distinct nuclear foci in wild-type flies. In *per*_*¹* mutants, TIM was diffusely distributed throughout the cytoplasm, with no detectable nuclear foci (Figure 2C), consistent with previous findings that PER/TIM heterodimerization is essential for TIM nuclear accumulation^19^.

### Timeless gene is localized close to the nuclear periphery in clock neurons

Next, we sought to investigate the spatial relationship between clock protein repressive foci and their target genes. To determine the subnuclear positioning of the *tim* gene, we first employed high-resolution DNA fluorescence in situ hybridization (DNA-FISH) using probes targeting a ∼10 kb region within the *tim* gene body. Fly brains were dissected at different circadian time points, and following tissue permeabilization, probe hybridization, and signal amplification, samples were imaged using confocal microscopy. We found that the *tim* gene localized near the nuclear periphery in clock neurons during both the activation and repression phases, with an average distance of ∼0.5 μm from the nuclear envelope (Figure 3A, B). Consistent with the fact that homologous chromosomes are paired in all Drosophila somatic cells^20^, including clock neurons, we detected only a single *tim* FISH signal per nucleus.

**Figure 3.**
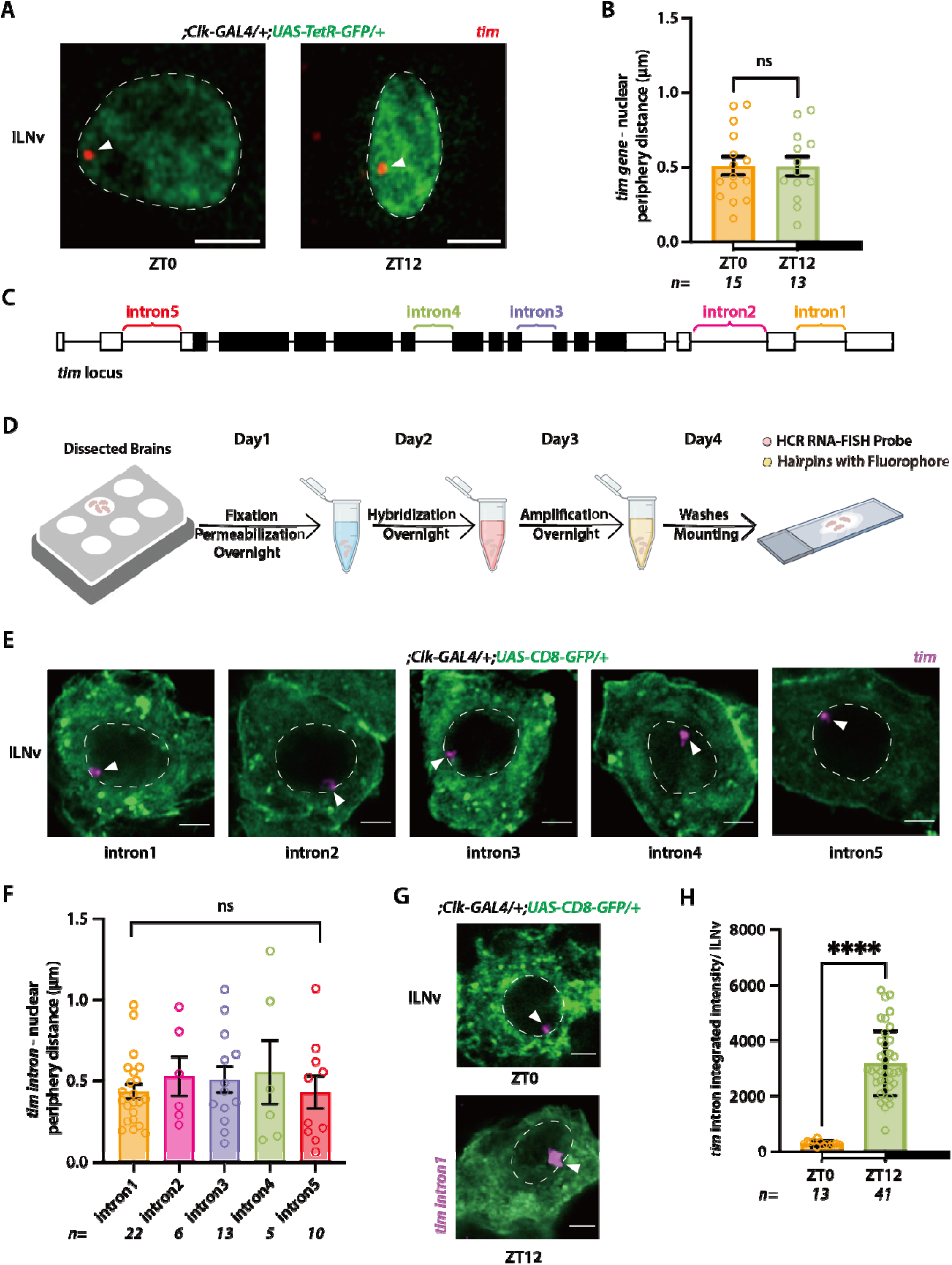
The *tim* gene is localized close to the nuclear periphery in clock neurons. (A) Representative DNA-FISH images of *tim* gene (red) in l-LNv neurons at ZT0 and ZT12. Scale bars-2 μm.(B) Quantification of the distance between *tim* gene loci and the nuclear envelope. (C) Schematic of the *tim* gene with introns indicated. Intron-specific probes were designed for HCR RNA-FISH. (D) Workflow of HCR RNA-FISH labeling. (E) Representative HCR RNA-FISH images of individual *tim* introns (magenta) in l-LNv neurons with cytoplasm labeled with GFP (green). Each intron signal corresponds to a transcription site, marking the location of the *tim* gene. Scale bars-2µm. (F) Quantification of the distance between *tim* introns and the nuclear envelope. (G) Representative HCR RNA-FISH images of *tim intron1* (red) in l-LNv neurons at ZT0 and ZT12. Scale bars-2µm. (H) Quantification of *tim intron1* fluorescence intensity at ZT0, ZT12. The statistical test used was a Kruskal-Wallis test (B, F, and H). ****P < 0.0001. Individual data points, mean, and SEM are shown. ‘n’ refers to the number of neurons.

We next attempted to combine DNA-FISH with protein imaging to simultaneously visualize PER condensates and *tim* gene loci. However, despite repeated optimization, PER fluorescence was completely lost during the high-temperature denaturation steps required for DNA-FISH, likely due to disruption of the fluorescent protein structure. To overcome this limitation, we implemented intron RNA-FISH using hybridization chain reaction (HCR-FISH) with probes targeting intronic regions of timeless. Because intron probes hybridize to unspliced nascent RNA at the gene locus^21^ as most introns are co-transcriptionally spliced, this approach provides a precise proxy for the genomic location of transcriptionally active genes. We also developed a combined protein imaging and RNA-FISH protocol that preserves protein fluorescence (see Methods).

First, we designed probes targeting multiple intronic regions of the *tim* gene and performed HCR-FISH on brains dissected at ZT0 and ZT12 (Figure 3C, D). The brightest intron signals corresponded to nascent RNA at active transcription sites, marking the gene’s physical location. Similar to the DNA-FISH results, *tim* transcription sites were consistently positioned near the nuclear periphery, with an average distance of less than 0.5 μm from the nuclear envelope (Figure 3E, F). Transcriptional activity was higher at ZT12 than at ZT0, as indicated by the increased frequency and intensity of intron signals (Figure 3G, H). This rhythmic transcriptional pattern persisted under constant darkness, although with slightly reduced amplitude (Figure S3).

### Clock gene loci are positioned adjacent to PER/TIM condensates at the nuclear periphery

Next, to probe the spatial relationship between PER/TIM foci and *tim* gene, we performed intron HCR-FISH in *per-mNeonGreen; Clk>CD4-tdTomato* flies. Notably, the PER fluorescence signal was well preserved, enabling simultaneous visualization of PER protein and *tim* gene loci within the same l-LNV clock neurons. We found that *tim* loci were positioned adjacent to PER condensates (Figure 4A, B), indicating a close spatial relationship between the PER foci and their target gene during the repression phase. As a negative control, we examined the shaker gene, which encodes a voltage-gated potassium channel involved in membrane repolarization and neurotransmitter release and is not under direct circadian control. We observed no spatial association between *shaker* loci and PER foci (Figure 4A, B). To assess whether this spatial relationship extends to other clock-regulated genes, we examined the *pdm3* gene, which is rhythmically expressed in DN1 neurons^21^. Similar to *tim*, *pdm3* loci were found adjacent to PER protein foci in DN1 cells (Figure 4C, D), revealing a close spatial relationship between repressive protein foci and their genomic targets. Together, these findings suggest a new model in which circadian transcriptional repression is spatially organized at the nuclear periphery, where clock proteins are sequestered away from chromatin into repressive condensates to coordinate the repression of clock-regulated genes (Figure 4E).

**Figure 4.**
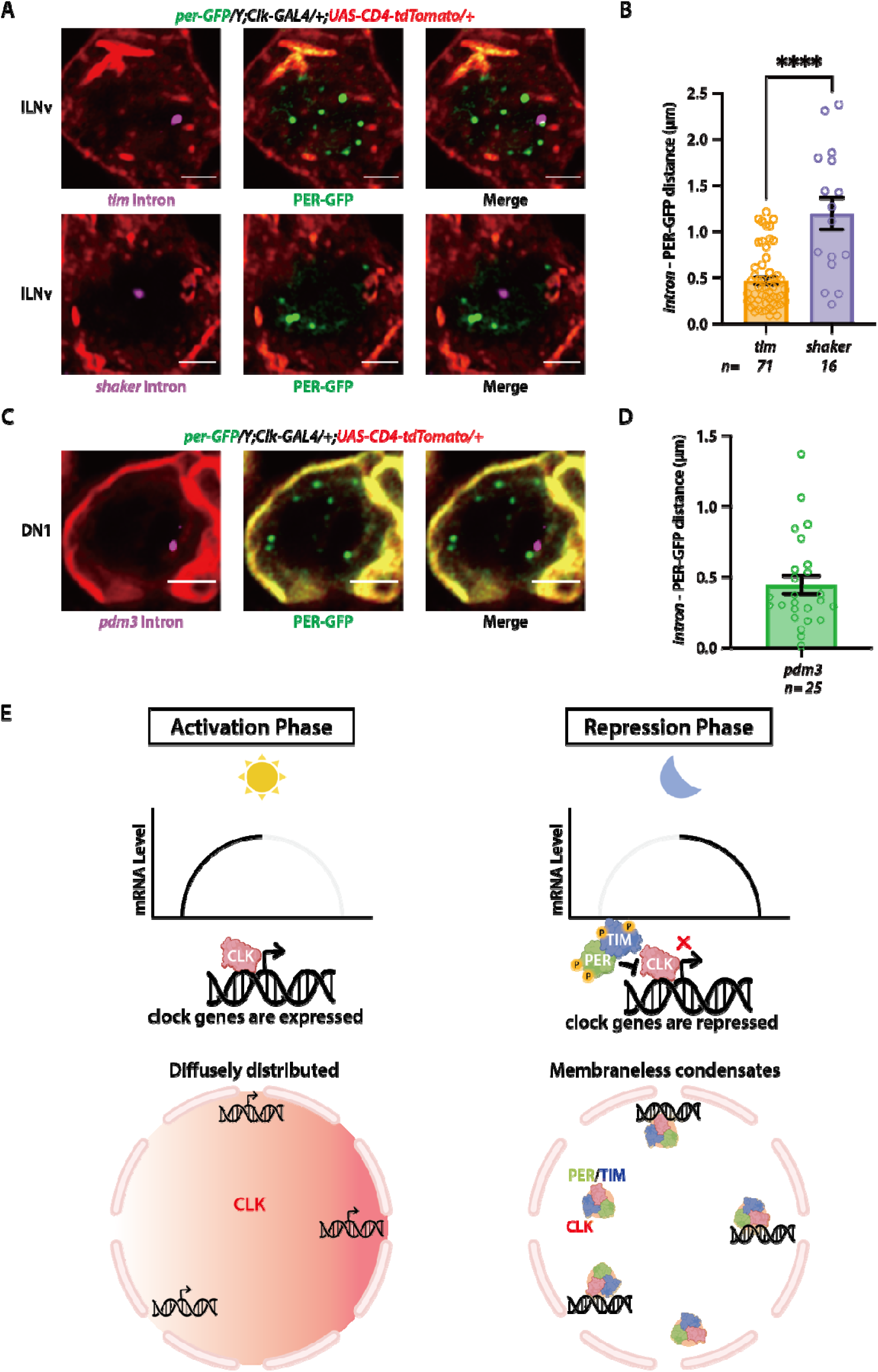
PER/TIM condensates were positioned adjacent to clock gene loci during the repression phase. (A) Representative images showing *tim* intron (magenta), *shaker* intron(magenta) and PER-GFP protein (green) in l-LNv neurons with cytoplasm labeled in tdTomato (red). Scale bars-2 μm. (B) Quantification of the distance between *tim, shaker* introns and PER-GFP foci in l-LNv neurons. (C) Representative images showing *pdm3* intron (magenta) and PER-GFP protein (green) in DN1 neurons. Scale bars-2 μm. (D) Quantification of the distance between *pdm3* introns and PER-GFP foci in DN1 neurons. The statistical test used was a Kruskal-Wallis test (B and D). ****P < 0.0001. Individual data points, mean, and SEM are shown. ‘n’ refers to the number of neurons.

Our findings provide a critical spatial dimension to the two-step biochemical model of circadian repression established by previous chromatin immunoprecipitation studies. This model proposes that repression begins with an “on-DNA” phase, in which PER is recruited to chromatin-bound CLK/CYC, followed by a stable “off-DNA” phase, where CLK is released from DNA and sequestered in a PER-CLK complex^22^. However, the physical nature and subnuclear location of this “off-DNA” complex have remained unknown. Our live-imaging data now reveal that this “off-DNA” state is not a pool of freely diffusing proteins but instead corresponds to phase-separated repressive condensates at the nuclear periphery. These condensates contain PER, TIM, and the sequestered activator CLK (as shown in our previous study), thus linking the biochemical two-step model to a concrete spatial mechanism. We propose that the “off-DNA” transition is mediated by a sequestration-by-condensation process, in which CLK is physically withdrawn from chromatin and incorporated into PER/TIM repressive condensates (Figure 4E). This provides a high-resolution framework for how nuclear architecture orchestrates rhythmic gene repression.

Traditionally, the nuclear periphery has been viewed as a repressive environment^23^. Our findings challenge this view by showing that both transcriptional activation and repression of clock genes occur within this compartment across the circadian cycle. We propose that the nuclear periphery acts as a dynamic regulatory hub, where temporal changes in the local molecular environment, such as the assembly and disassembly of clock protein condensates, dictate whether genes are actively transcribed or repressed.

Based on our findings and recent cross-species observations of clock protein nuclear bodies^2, 6-11^, we propose that organisms have evolved a mechanism allowing clock proteins to undergo regulated phase transitions between diffuse (activation) and condensed (repression) states. This dynamic behavior enables the temporal organization of transcriptional repression over 24-hour cycles. The combination of protein condensate formation and gene positioning likely imparts robustness and precision, ensuring that circadian rhythms remain stable under environmental fluctuations. By concentrating key regulatory proteins, this spatial organization buffers against molecular noise and enforces a cooperative, switch-like transition into the repressed state. Together, these findings reveal a previously unrecognized layer of circadian regulation, driven by the spatial coordination between clock proteins and chromatin.

## MATERIALS AND METHODS

### Fly stocks

Flies were raised on standard cornmeal/yeast media and maintained at 25C under a 12h:12h LD schedule. The following flies used in the study were previously described or obtained from the Bloomington Stock Center: w1118 (BL-6326), Clk-GAL4^24^, *per^01^* ^25^, UAS-CD8GFP (BL-5137), UAS-CD4tdTom (BL-35837), *tim^01^* ^5^. Flies were entrained to Light-Dark (LD) cycles where they were exposed to 12-hour Light-Dark (LD) cycles for 5-7 days, followed by a shift to complete darkness (DD) for another 6–7 days. The initiation of the light phase is labeled as Zeitgeber Time (ZT) 0, and ZT12 signifies the beginning of the dark period. When referencing the times in the continuous darkness phase, we use Circadian Time (CT), with CT0 marking the time when lights would have been turned on and CT12 the time when lights would have been turned off. The labels DD1 and DD2 represent the first and second days of complete darkness, respectively.

### Endogenous fluorescence tagging using CRISPR/Cas9

*Tim-mNeonGreen flies*: To generate *tim-mNeonGreen* fly, we used the protocol described in our previous paper[JOVE]. Donor plasmid is directly ordered from Genscript. EcoRI, XbaI-linearized pBluescript sk(-) plasmid backbone was mixed with 5’ arm, mNeonGreen fragment, and 3’ arm fragments to synthesize sgRNA plasmids.

Two pairs of gRNA primers are used:

tim-gRNA1-sense: cttcACTTGAGCGAACCGAGATCC,

tim-gRNA1-antisense:aaacGGATCTCGGTTCGCTCAAGT,

tim-gRNA2-sense: cttcGGTGGGCCTCACTCAAAACT,

tim-gRNA2-antisense: aaacAGTTTTGAGTGAGGCCCACC.

Golden Gate Assembly protocol was applied with type IIS endonuclease Esp3I (NEB). The product was transformed into competent E. coli cells (NEB, C2987H) and plated on LB-Amp plates. Positive colonies were purified (QIAGEN, Plasmid Mini Prep Kit), confirmed by Sanger sequencing and prepared for injection (QIAGEN, Plasmid Midi Kit). After injected embryos developed into adult flies, these adult flies were individually crossed to sp/cyo;ubx/hu male or virgin female flies. Positive progenies with the presence of the insertion were screened by single-leg PCR. We backcrossed the flies that had the insertion to sp/cyo;ubx/hu flies and then homozygosed their progeny. Homozygous stocks were genotyped and Sanger-sequenced.

*Per-mScarlet* flies: We used the same protocol as described as above. The sgRNA sequence information can be found in our previous paper^2^.

### Live imaging protocol

Flies were entrained to Light-Dark (LD) cycles with lights on for 12 hours and off for 12 hours for 5-7 days, and then released into complete darkness (DD) for 6-7 more days. All flies used for live imaging experiments were placed in density-controlled food vials (3 females and 1 male) and entrained for 5-7 days in incubators. We performed all our live imaging experiments on 5-7 day old male or female flies, and did not notice any differences in our experimental results. We used the GAL4/UAS system to express transgenes in clock neurons in the brain.

For imaging of live clock neurons, 3-4 brains were dissected in PBS medium in less than 5 minutes under low light conditions. A punched double-sided tape was used as a spacer on the slides to prevent flattening of the brains. The brains were overlaid with a small amount of ProlongTM Glass Antifade mounting medium (ThermoFisher Scientific, P36982) and covered using a coverslip. We imaged our samples using a Zeiss LSM800 laser scanning confocal microscope with AiryScan superresolution module (125 nm lateral and 350 nm axial resolution). We have acquired our images using a 63x Plan-Apochromat Oil (N.A. 1.4) objective and 405, 488, and 561 nm laser lines. We collected Z-stack (each Z-slice is about 250 nm) or time-lapse image series of individual clock neurons and images were analyzed using Zeiss ZEN software and ImageJ.

### RNA fluorescence in situ hybridization (RNA-FISH)

We followed the following protocol for Hybridization Chain Reaction Fluorescence In Situ Hybridization (HCR-FISH). In brief, brains were dissected in PBS buffer, fixed for 20min at room temperature in 4% PFA, washed with PBS buffer, and put in 70% ethanol overnight at 4C for permeabilization. Next day, brains were washed in FISH wash buffer (2xSSC, 10% formamide, 2mM RVC, 0.2% Tween-20) for 5min at room temperature, and subsequently hybridized at 37C overnight for a minimum of 16-hours (2xSSC, 10% formamide, 10% dextran sulfate, 2mM RVC, 0.2% Tween-20, 5nM probe). Hybridized brains were washed in FISH wash buffer two times for a total of 1-hour wash at 37C to remove unbound probes and subsequently mounted in Prolong Glass mountant (smFISH) or proceed with amplification (HCR-FISH) overnight for a minimum of 16-hours (2xSSC, 10% dextran sulfate, 2mM RVC, 0.2% Tween-20, 60nM of each amplifier hairpin). Amplified brains were washed in 2xSSC + 0.2% Tween-20 two times for a total of 1-hour before mounting. Probes and amplifiers were purchased from Molecular Instruments (HCR-FISH).

### Combined protein and HCR-FISH imaging

To perform Hybridization Chain Reaction Fluorescence In Situ Hybridization (HCR-FISH) in whole-mount Drosophila brains, we followed the same protocol in the previous paper[plos genetics]. Probes and amplifier hairpins were synthesized by Molecular Instruments. Notably, we found an extra 10 minutes incubation in ProlongTM Glass Antifade mounting medium (ThermoFisher Scientific, P36982) at room temperature before fixation can help preserve fluorescent protein signals. For all experiments, we housed the flies in density-regulated food vials, each containing 8 females and 2 males. They were acclimated in incubators for 5–7 days. All imaging tests were conducted on male or female flies aged between 5–7 days and no differences were observed in the outcomes between genders. Specific fly genotypes and Zeitgeber Time (ZT) details are provided in the accompanying Figure legends. We employed the GAL4/UAS system to express transgenes in the brain’s clock neurons. We captured our images using the Zeiss LSM800 confocal microscope, equipped with an AiryScan super-resolution module that offers 125 nm lateral and 350 nm axial resolution. Imaging was done utilizing a 63x Plan-Apochromat Oil (N.A. 1.4) objective, with laser lines of 405, 488, and 561 nm. Our imaging sessions included collecting Z-stack series (with approximately 250 nm per Z-slice) of distinct clock neurons. Following image acquisition, we imported the CZI files into ImageJ, ensuring we maintained a lossless 16-bit resolution for each channel, using the Bio-Formats Importer to process them as composite images. Subsequently, we manually delineated regions of interest (ROIs) on the channel, fine-tuning the white values to optimize visualization. To guarantee an unbiased fluorescence analysis, we remained consistent with our visualization settings across all images, keeping them specific to each fly line and neuron type.

### Locomotor activity and Rhythmicity analysis

Individual adult male flies (3-5 days old) were placed in glass capillary tubes (∼4 mm inner diameter, 5 cm in length) containing 2% agar and 4% sucrose food, which were then loaded into TriKinetics DAM2 Drosophila Activity Monitors (Waltham, MA, USA) for locomotor activity recordings. Flies were entrained to Light-Dark (LD) cycles with lights on for 12 hours and off for 12 hours for 5-7 days, followed by complete darkness (DD) for six more days. The DAM monitors are equipped with infrared sensors and they record infrared beam breaks when the flies cross the middle of the glass tube. The monitors with flies were placed in the incubators and the beam crossing counts from each monitor were recorded on a computer. The Beam crossing counts were placed into 30-minute bins for time-series analysis of locomotor activity. Averaged population activity profiles under Light-Dark cycles and constant conditions were generated using a commercially available software ClockLab (Actimetrics) and public domain R Rethomics software package. Briefly, activity levels were normalized for each fly, such that the average number of beam crossings in each day (48 bins) is equal to 1. Next, the population average of normalized activity was determined and the results are displayed as normalized activity plots in the figures. We used activity counts of individual flies under complete darkness conditions following LD cycles to analyze rhythmicity and to determine the free-running period of the circadian clock. Rhythmicity and free-running period of individual flies were determined by a chi-square periodogram analysis with a confidence level of 0.001 using the ClockLab software. The “Power” and “Significance” values generated from the chi-square analysis were used to calculate “Rhythmic Power” as a measure of the strength of each rhythm.

### Data Analysis

*Analysis of foci fluorescence intensity*: To guarantee an unbiased fluorescence analysis, we remained consistent with our visualization settings across all images, keeping them specific to each fly line and neuron type. First, we selected a single Z-plane with the largest foci count or brightest HCR-FISH dot for each cell. Each focus was annotated with a polygon region of interest (ROI) for analysis. The mean pixel brightness (arbitrary unit, a.u.) and geometric area (!“#) of the ROI were acquired using built in functions of ImageJ software. The integrated intensity of the focus is obtained from a simple multiplication of pixel brightness and geometric area, yielding numeric value with unit a.u.. Next, we subtracted the background fluorescence from the integrated intensity of the ROI region (ImageJ). Background fluorescence per pixel was estimated by the mean intensity of a background selection on the same plane.

*Analysis of distance between TIM protein foci or tim intron and the nuclear periphery*: First, we selected the center Z-plane for each cell where the nuclear envelope is clearly visible. Next, foci were annotated with elliptical ROIs. In the same center Z-plane, nuclear periphery was annotated with closed polygon ROIs. Finally, we calculated the shortest distance from the centroid of each foci to the respective closed polygon.

*Analysis of foci count and percentage of cells with foci*: We used appropriate white values for visualization of our acquired images. We manually annotated the foci if they are clearly distinct from the background. Both analyses were performed independently by manually counting foci across all Z-planes for each cell.

## ACKNOWLEDGEMENTS

We thank members of the Yadlapalli laboratory for helpful discussions and Yangbo Xiao for help with the design of PER-mScarlet CRISPR-edited flies and Christopher Wilson for help with initial set of experiments. We thank the Bloomington Drosophila Stock Center for providing fly strains.

This work was supported by National Institutes of Health NIGMS R35GM133737 grant and NINDS RO1NS140761 grant to S.Y., the Alfred P. Sloan fellowship, the McKnight scholar grant, and the Chan Zuckerberg collaborative pairs grant to S.Y., the Rackham International Student Fellowship to Q.Q., and the Rackham Predoctoral Fellowship to Y.Y..

## AUTHOR CONTRIBUTIONS

Q.C. performed protein and FISH imaging experiments and analyzed the data. Y.Y. performed the CRISPR experiments to generate TIMELESS-MneonGreen knock-in flies. D.C., and S.Y. performed live imaging experiments. Q.C. and Y.Y. wrote the manuscript with input from all authors.

## CONFLICTS OF INTEREST

The authors declare no financial conflict of interest.

---

**Supplementary Figure 1.**
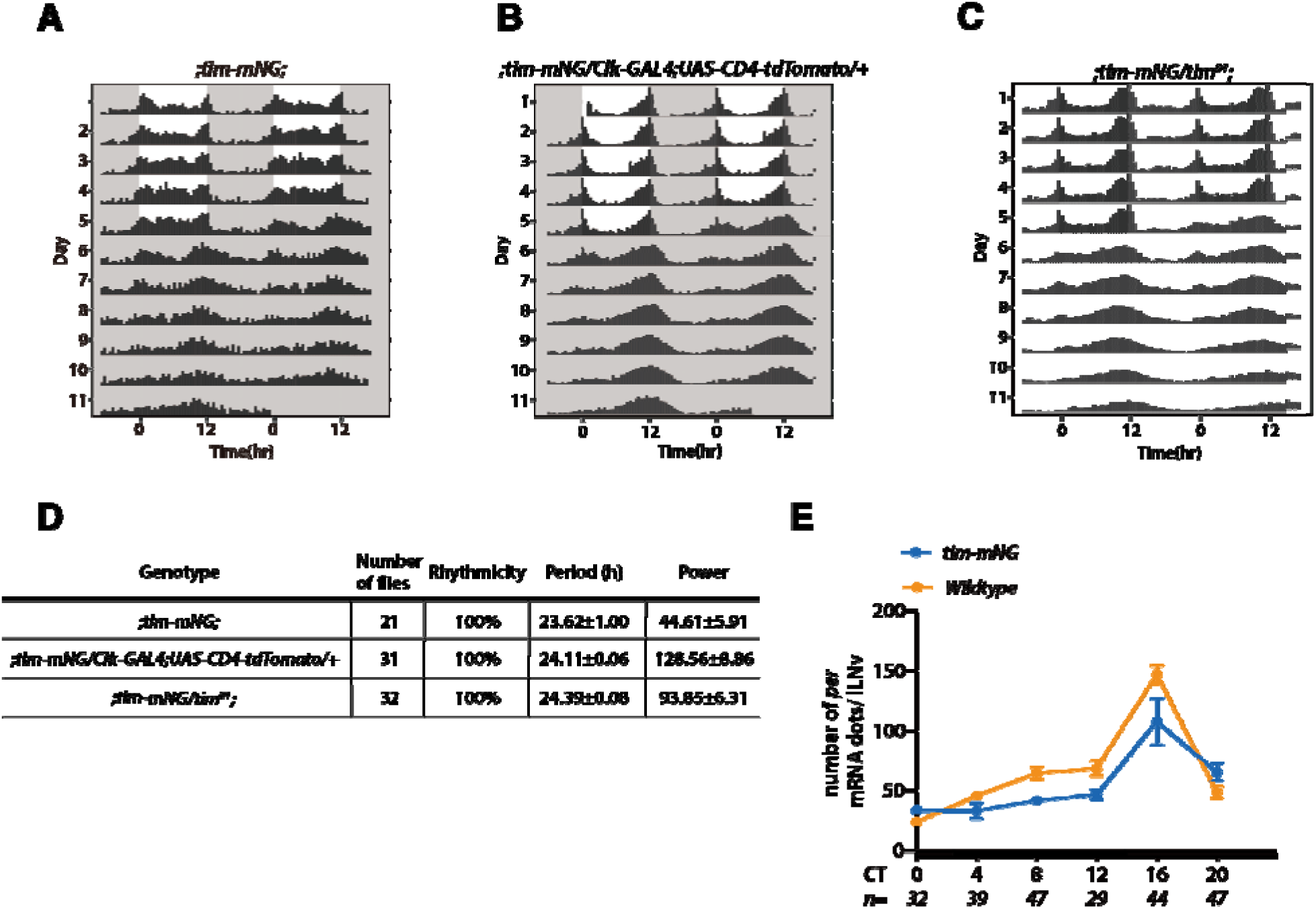
*tim-mNeonGreen* flies exhibit normal circadian behavior and robust per mRNA oscillations. (A-C) Representative locomotor activity actograms of different genotypes under constant darkness (DD) conditions. (A) *tim-mNG* flies; (B) *tim-mNG/Clk-GAL4; UAS-CD4-tdTomato/+* flies; (C) *tim-mNG/tim^01^* flies. Each row represents two consecutive days of activity; shaded areas indicate night. (D) Summary table of rhythmicity, period length, and power across indicated genotypes. (E) Quantification of *per* mRNA levels by single molecule FISH (smFISH) assay in *tim-mNG* (blue) and *y,sc,v* control (orange) flies across circadian cycle. Both genotypes exhibited similar daily oscillations in *per* transcript abundance. ‘n’ refers to the number of neurons.

**Supplementary Figure 2.**
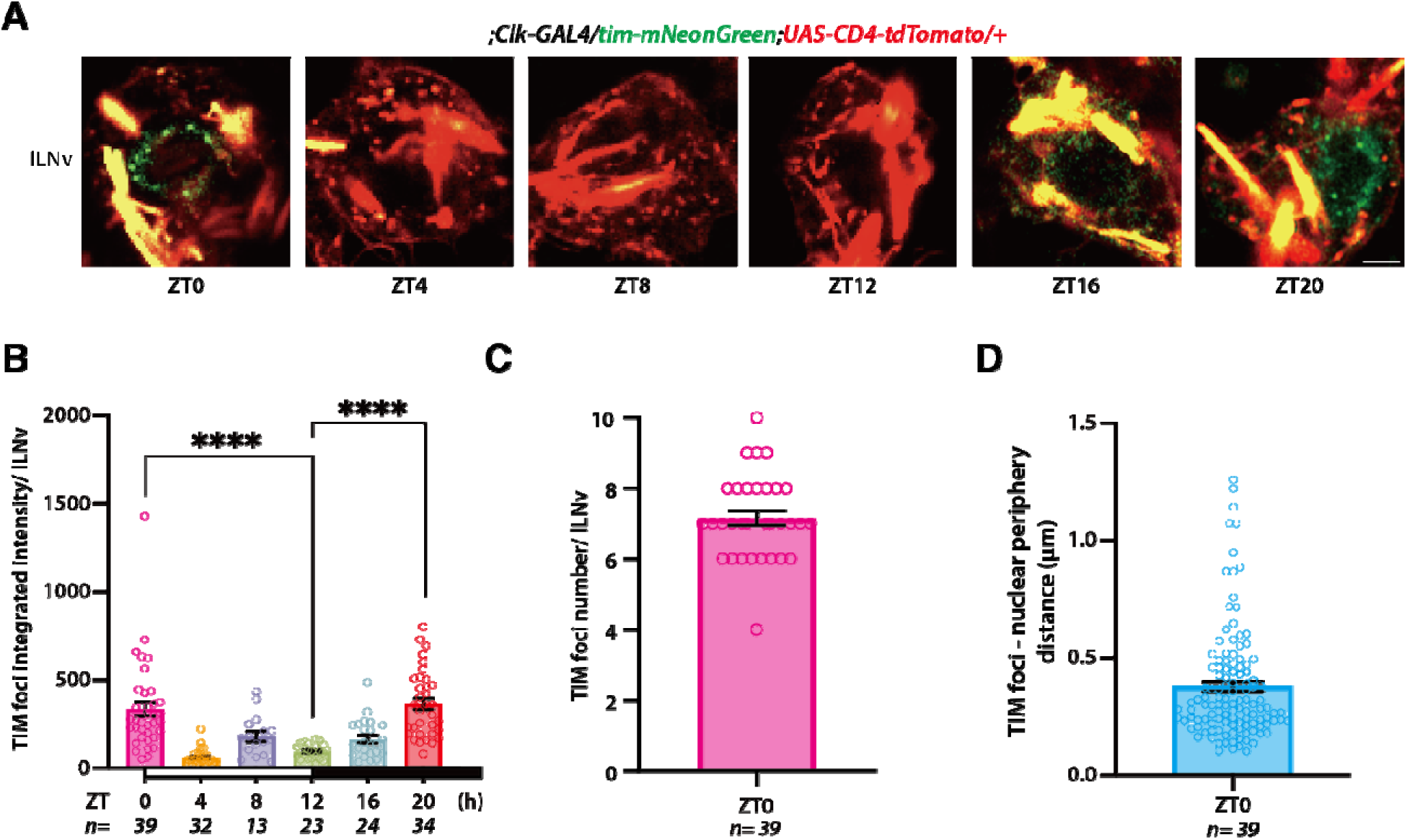
TIM is organized into nuclear foci in l-LNvs during the repression phase. (A) Representative images of TIM-mNeonGreen (green) in l-LNv neurons at different zeitgeber times (ZT0, ZT4, ZT8, ZT12, ZT16, ZT20). CD4tdTom (red) is used to label the membranes in the cytoplasm. (B) Quantification of TIM-mNeonGreen fluorescence intensity per l-LNv across circadian cycle. (C) Quantification of the number of TIM-mNeonGreen foci per l-LNv at ZT0. (D) Distance of TIM-mNeonGreen foci to the nuclear envelope at ZT0. The statistical test used was a Kruskal-Wallis test (B, C and D). ****P < 0.0001. Individual data points, mean, and SEM are shown. ‘n’ refers to the number of neurons.

**Supplementary Figure 3.**
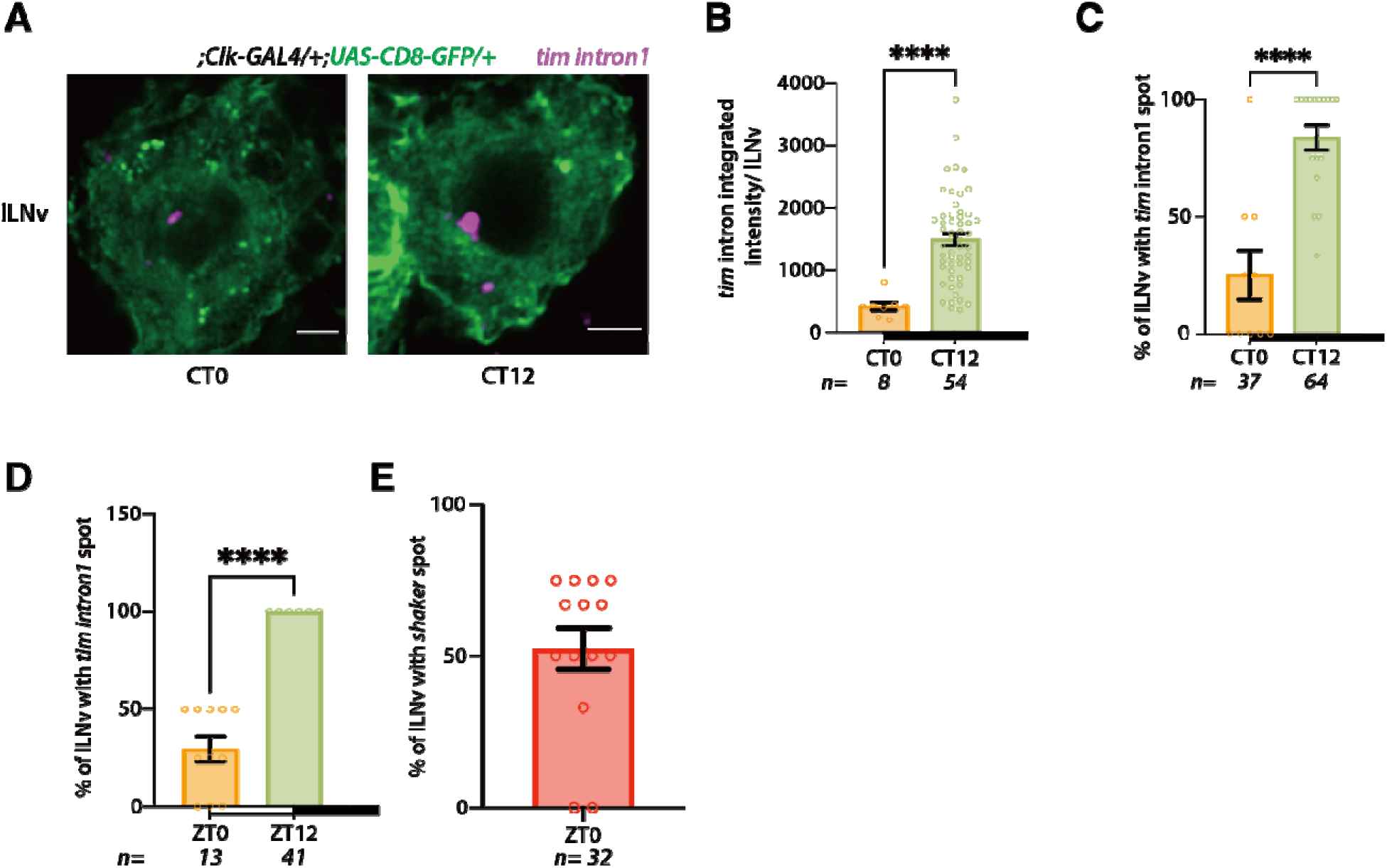
*tim* transcriptional activity is rhythmic in the circadian cycle. (A) Representative HCR RNA-FISH images of *tim* intron1 (magenta) in l-LNv neurons at CT0 and CT12 on the first day of constant darkness (DD). Flies were entrained to light-dark (LD) cycles for 5 days before release into DD. CT refers to circadian time. Scale bars-2 μm. (B) Quantification of *tim* intron1 spot intensity at two timepoints across the circadian cycle. (C) Percentage of l-LNv neurons with detectable *tim* intron1 spot at CT0 and CT12. (D) Percentage of l-LNv neurons with detectable *tim* intron1 spot at ZT0 and ZT12. (E) Percentage of l-LNv neurons with detectable *shaker* intron spot, which showed no circadian modulation. The statistical test used was a Kruskal-Wallis test (B, C and D). ****P < 0.0001. Individual data points, mean, and SEM are shown. ‘n’ refers to the number of neurons.

## Notes

### Competing Interest Statement

The authors have declared no competing interest.

